# Hemispherotomy: a cortical island of sleep-like activity in awake humans

**DOI:** 10.1101/2025.02.10.637383

**Authors:** Michele A. Colombo, Jacopo Favaro, Ezequiel Mikulan, Andrea Pigorini, Flavia M. Zauli, Ivana Sartori, Piergiorgio d’Orio, Laura Castana, Irene Toldo, Stefano Sartori, Simone Sarasso, Tim Bayne, Anil K. Seth, Marcello Massimini

## Abstract

Hemispherotomy is a neurosurgical procedure for treating refractory epilepsy, which entails disconnecting a significant portion of the cortex, potentially encompassing an entire hemisphere, from its cortical and subcortical connections. While this intervention prevents the spread of seizures, it raises important questions. Given the complete isolation from sensory-motor pathways, it remains unclear whether the disconnected cortex retains any form of inaccessible awareness. More broadly, the activity patterns that large portions of the deafferented cortex can sustain in awake humans remain poorly understood.

We address these questions by exploring for the first time the electrophysiological state of the isolated cortex before and after surgery in ten awake pediatric patients. Post-surgery, the isolated cortex exhibited prominent slow oscillations (<2 Hz) and a broad-band shift in power spectral density from high to low frequencies. This resulted in a marked decrease of the spectral exponent, a validated consciousness marker, indicating broad-band slowing characteristic of unconscious states.

When compared with a reference pediatric sample across the sleep-wake cycle, the spectral exponent of the contralateral cortex aligned with wakefulness, whereas that of the isolated cortex was consistent with deep NREM sleep. However, spindles did not emerge in the isolated cortex due to the lack of subcortical inputs, constituting a fundamental difference from physiological sleep.

These findings demonstrate a unihemispheric sleep-like state during wakefulness, challenging the possibility that hemispherotomy might lead to inaccessible “islands of awareness.” Moreover, the persistence of sleep-like patterns years after disconnection provides unique insights into the electrophysiological effects of disconnections in the human brain.

## Introduction

Hemispherotomy is a surgical procedure used to treat severe cases of refractory epilepsy in children (Griessenauer et al., 2015; Marras et al., 2010). The procedure aims at maximal disconnection of the pathological cortex from the rest of the brain by severing its white matter connections with brainstem, basal ganglia, thalamus and contralateral hemisphere. Typically, the extent of the cortical disconnection in hemispherotomy ranges from one lobe to an entire hemisphere (Marras et al., 2010; Rizzi et al., 2019). The disconnected/isolated cortex, completely devoid of communication with the external environment (except for cortical afferents from the olfactory bulb, in some cases) is left inside the cranial cavity with preserved vascular supply (de Ribaupierre and Delalande, 2008). This neurosurgical procedure is effective in blocking the spread of epileptic seizures from the pathological cortex to the rest of the brain, allowing patients to lead satisfying lives (Marras et al., 2010; Delalande et al. 2007; de Palma et al., 2019). It is clear that the intact contralateral cortex (which remains connected to the body and the external world) continues to support wakeful consciousness after surgery, as patients recover (at least) partial sensory-motor functions, cognitive abilities and the capacity to report their internal states (Marras et al., 2010; Devlin et al., 2003). However, the disconnected cortex, isolated from sensory and motor pathways, cannot be evaluated behaviorally, leaving open the question of whether it retains internal states consistent with some form of awareness (Bayne, Seth and Massimini, 2020; Gauvry and Rüber, 2024). More broadly, the activity patterns that extensive regions of the human brain can sustain following massive deafferentation remain unknown.

The available evidence is drawn from preliminary fMRI studies. A first study, involving two pediatric participants, revealed preserved lateralized connectivity in resting state networks within the disconnected hemisphere, despite decreased regional cerebral blood flow (Blauwblomme et al., 2020). A classifier aligned the pattern in the disconnected cortex with that of Minimally Conscious State (MCS) patients, indicative of a state of diminished consciousness. A subsequent study in a larger cohort found preserved functional connectivity in all resting-state networks, including the default mode network (DMN), in the disconnected hemisphere, albeit with reduced between-network segregation (Rüber et al. 2024). This finding was interpreted as indicative of a potential ‘island of awareness’. However, inferring the internal state of the isolated cortex solely based on fMRI activity remains challenging. Indeed, functional connectivity within resting-state networks is observed not only during wakefulness but also in states such as coma, general anesthesia, and deep Non-Rapid Eye Movement (NREM) sleep (Boly et al., 2008; Horovitz et al., 2008; Vincent et al., 2007; Akeju et al., 2014). Adding further complexity, brain injuries and structural disconnections have been shown to produce mixed states, where sleep-like activity dominated by EEG slow waves intrudes upon the awake brain (Massimini et al., 2024).

One way to clarify this complex landscape would be to assess the regional electrophysiological states of the brain of awake subjects both before and after hemispherotomy. To date, the only EEG study conducted on hemispherotomy patients was performed under general anesthesia, which precludes addressing this question (Hawasli et al., 2017).

Here, we fill this gap by studying for the first time regional EEG patterns during wakefulness in ten pediatric participants who had undergone a complete cortical disconnection, of either an entire hemisphere (complete hemispherotomy in 7 cases) or a large portion of it (temporo-parieto-occipital disconnection in 3 cases).

After hemispherotomy we observed the emergence of prominent slow waves over the disconnected cortex, but not over the contralateral hemisphere. These slow waves are reminiscent of those observed in deep sleep (Merica et al., 1997; De Gennaro et al., 2001; Ogilvie, 2001), during anesthetic-induced loss of consciousness (reviewed in Brown et al., 2010) and in disorders of consciousness (Lechinger et al., 2013; Chennu et al., 2014; Sitt et al., 2014; Estraneo et al., 2016; Schiff, 2016; Comanducci et al., 2020; Colombo et al., 2023). We accordingly quantified slow wave activity by narrow band spectral analysis within the Slow-Delta band (0.5-2 Hz), after excluding all visible epileptiform activity. To evaluate the possibility that the isolated hemisphere may retain some level of residual consciousness, we then estimated the EEG spectral exponent. This measure, reflecting the 1/f-like decay of the Power Spectral Density (PSD), provides a comprehensive quantification of broad-band electrophysiological slowing, and has been previously validated as a reliable index of consciousness across various physiological (Shen et al., 2003; Miskovic et al., 2019; Bódizs et al., 2021; Zilio et al., 2021; Schneider et al., 2022; Horváth et al., 2022; Alnes et al., 2023; Rosenblum et al., 2024; Bódizs et al., 2024) pharmacological (Zilio et al., 2021; Colombo et al., 2019; Maschke et al., 2023a; Maschke et al., 2023b; Zilio et al., 2023; Casey et al., 2023) and pathological conditions (Colombo et al., 2023; Maschke et al., 2023a; Zilio et al., 2023), including states of disconnected consciousness, in the form of ketamine-induced dreaming and the locked-in syndrome. Importantly, the spectral exponent has been extensively tested in large pediatric samples during wakefulness (Favaro et al., 2023; Cellier et al., 2021; McSweeney et al., 2023; Schaworonkow and Voytek, 2021) and NREM sleep (Schneider et al., 2022; Favaro et al., 2023).

Finally, to further characterize the electrophysiological state of the cortex before and after hemispherotomy we directly compared it to the patterns recorded in a reference sample of healthy controls in the same age range during both wakefulness and NREM sleep (Favaro et al., 2023).

## Materials and methods

### Patients receiving disconnective surgery

We retrospectively selected our participants starting from a large sample of pediatric patients referred from 2005 to 2020 to the Munari Center for Epilepsy Surgery of Niguarda Hospital in Milan. Ten participants were recruited on the basis of the following inclusion criteria: diagnosis of focal drug-resistant epilepsy with structural etiology on a malformative basis or as an outcome of perinatal hypoxic-ischemic insult; patient undergoing complete cortical disconnection of an hemisphere or of a large portion of it (at least one lobe); post-intervention outcome classified in class I according to the Engel Surgical Outcome Scale; availability of an EEG within 12 months prior to surgery and at least six months after surgery; age at pre-surgery EEG recording between 2 and 17 years; presence of at least 3 min of quiet wakefulness free from artifacts and interictal epileptiform discharges.

During the EEG the participants, lying on a bed or in the arms of the parent, did not perform any particular task, except alternating periods of dozens of seconds with eyes open and with eyes closed, following a constant protocol across participants. Since we were not interested in the neural basis of specific contents of consciousness, we considered these epochs jointly (as in Favaro et al., 2023). We excluded from the analysis all epochs in which the clinicians performed activation tests (intermittent light stimulation and hyperpnea). When necessary, younger children watched cartoons in order to improve compliance and reduce motion artifacts. Standard EEG recordings, performed for routine clinical purposes, were used for research purposes after acquiring a written informed consent from the participants’ parents. The study was conducted in accordance with the Declaration of Helsinki and the institutional guidelines and the protocol was approved by the Ethics Committee of Niguarda Hospital, Milan, Italy.)

### Data acquisition and pre-processing

Each EEG recording was obtained by 19 scalp electrodes (cup in chlorinated silver) positioned according to the international 10–20 system and a 32-channel amplifier (Neurofax EEG-1200, Nihon Kohden Corporation). In acquisition, an additional electrode located between the Fp1 and Fp2 electrodes was used as a reference; the impedances of all the electrodes were kept below 5 kΩ for the entire duration of the recording. The signal was filtered during acquisition (high-pass filter 0.016 Hz; low-pass filter 300 Hz) and sampled at 500 Hz. The raw data was imported and analyzed with custom MATLAB code (MATLAB 9.7.0 R2019b, The MathWorks Inc., Natick, MA, USA). The signal was filtered with a 5th order Butterworth high-pass filter at 0.1 Hz (effectively retaining Slow-Delta frequency activity between 0.5 Hz and 2 Hz), with a 3rd order Butterworth low-pass filter at 45 Hz and with a 50 Hz Notch filter. Epochs characterized by artifacts and interictal epileptiform discharges (spikes, polyspikes, sharp waves, spike/polyspike-wave complexes) were visually selected and excluded by a trained pediatric neurologist expert in EEG **(Supplementary Figure 1).** Due to the careful application of electrodes in our low-density set-up, recordings from all electrodes were deemed of sufficiently good quality. Electrodes were first re-referenced to the common average, to ease the topographical inspection of the components from a subsequent blind source separation. Specifically, to minimize the influence of electromyographic (EMG) and electro-oculographic (EOG) activity, Independent Component Analysis was performed on a copy of the data, filtered with a narrower high-pass filter (5th order Butterworth at 0.5 Hz) to better separate artifactual from neurophysiological components. The different components of the signal were visually inspected on the basis of their time-series, topographies and PSD. Clearly identifiable components resembling EMG and EOG activity were marked for rejection. The unmixing weights (estimated from the data filtered with a 0.5 Hz high-pass) were applied to the original data (i.e. that filtered with a 0.1 Hz high-pass), and the retained components were back-projected to the scalp, thus effectively removing artifactual influences from the scalp electrodes signal (Winkler et al., 2015).

Finally, to observe lateralized effects with high spatial specificity under our low-density EEG montage, bipolar derivations between all pairs of nearest neighboring electrodes (excluding the midline) were considered for subsequent analyses. The common average reference was used exclusively for ICA preprocessing and for topographic visual inspection and display (**Figure 1B, center).**

**Figure 1.**
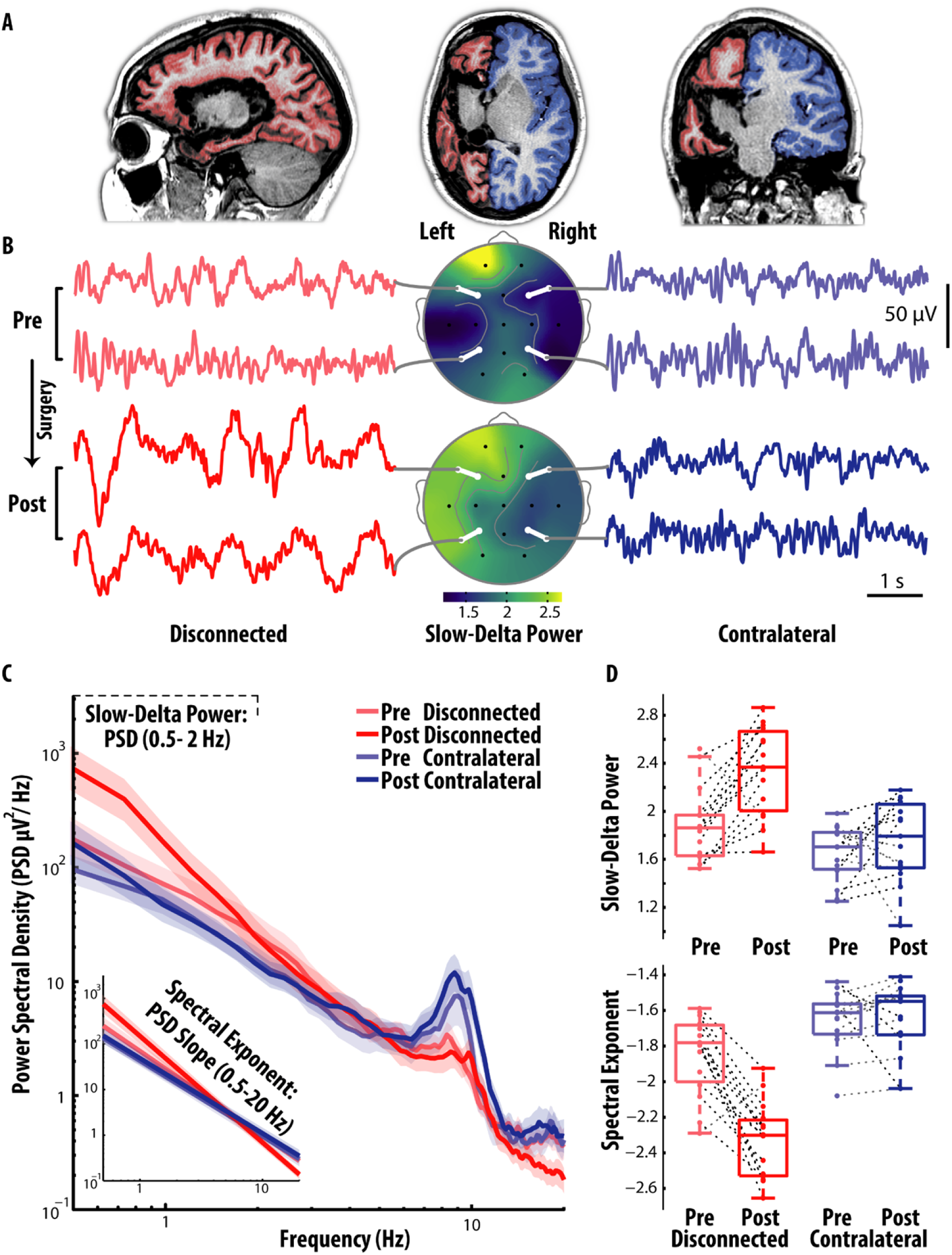
Representative narrow- and broad-band spectral changes after hemispherotomy , displaying marked EEG slowing of the isolated cortex. Cortical isolation of the left hemisphere in a 11 y.o. representative patient. **(A)** The anatomical MRI 6 months after surgery shows that all interhemispheric and subcortical connections were resected (sagittal, axial and coronal panels are shown from left to right). The cortical surface is highlighted in red for the disconnected cortex (in blue for the contralateral). **(B)** EEG recordings were performed 8 months before surgery (at 11 y.o.), and 12 months after. Short EEG segments (6 s) from the isolated cortex (F7-F3 and T5-P3 bipolar derivations, left panel) and the contralateral homologue (F8-F4 and T6-P4, right panel) are shown, pre-surgery (above) and post-surgery (below). In the middle panel, the topographies of Slow-Delta power, indexing the power spectral density (PSD) of low-frequency activity (0.5-2 Hz, hereby estimated under average reference for display purposes), show a marked inter-hemispheric asymmetry post-surgery. **(C)** The EEG PSD, averaged by geometric mean across intrahemispheric bipolar derivations, reveal an increase of slow frequency activity (< 2 Hz) and a broad-band steepening of the PSD over the disconnected hemisphere post-surgery, which is evidenced in the inset graph by the the linear fit of the PSD under double logarithmic axes (power-law fit). The shaded area indicates the bootstrapped 95% confidence interval for the geometric mean across electrodes. **(D)** For all the neighboring intrahemispheric bipolar derivations, we display the values of Slow-Delta power (0.5-2 Hz), and of the spectral exponent, indexing the slope of the PSD decay (0.5-20 Hz).

### Estimation of slow delta power and spectral exponent

We estimated narrow-band EEG slowing by means of Slow-Delta power and broad-band EEG slowing by means of the spectral exponent. Both metrics were derived by the PSD, which was estimated using Welch’s method, with a 3 second Hanning window and a 50% overlap, following linear detrending of the time-series in each periodogram. Only periodograms that were entirely free from artifacts and from epileptiform activity were included. The estimation of low frequency activity (above 0.5 Hz) was free from artifacts induced by the filter (0.1 Hz high-pass, as previously mentioned).

We estimated Slow-Delta power as the mean absolute PSD between 0.5 and 2 Hz, subsequently transformed by log10.

The spectral exponent, on the other hand, indexes the slope of the overall PSD decay across frequencies, and thus characterizes the broad-band relative distribution between high and low frequencies (1/f-like). The neurophysiological 1/f-like distribution originates from aperiodic and quasi-periodic activity (Palva and Palva 2018; Gerster et al., 2022).

The exponent was estimated from a linear fit of the PSD under double logarithmic axes, after discarding frequency bins with peaks corresponding to periodic activity, i.e., bins with large positive residuals from a preliminary linear fit, and their contiguous bins with positive residuals (**Supplementary Material, section 2, 3, Supplementary Figure 2**, see Colombo et al., 2019). The Matlab code for PSD fitting, and an equivalent Python code translation, is available online https://github.com/milecombo/spectralExponent/blob/master/README.md.

Here, we estimated the spectral exponent over the 0.5–20 Hz range, so as to include the Slow-Delta range, while excluding higher frequencies which are prone to muscular contamination in children. A similar fitting range, 1-20 Hz, previously used in Favaro et al., 2023, was also investigated in our patients’ sample for completeness (**Supplementary Material, section 4; Supplementary Figure 3**).

Subsequently, the median value of each feature (Slow-Delta power and spectral exponent) was taken across the lateralized bipolar derivations placed over the isolated cortex (15 derivations in the case of a hemispherotomy and 6 for a temporo-parieto-occipital disconnection) and the corresponding homologous bipolar derivations placed over the contralateral cortex (see additional analysis in **Supplementary Material, section 5**).

### Statistical comparisons: spatial inter-hemispheric differences, and temporal pre-post differences

We estimated narrow-band and broad-band EEG slowing, by means of Slow-Delta power and of the spectral exponent, before and after surgical disconnection, in both the disconnected and in the homologous contralateral cortex. For each of the two EEG features, we thereby assessed whether EEG slowing differed between homologous cortices (spatial effect), between the recording sessions (pre vs post-surgery, temporal effect), and if it differently changed in the two cortices following surgery (spatio-temporal interaction effect), by means of an ANOVA on a mixed effects model (**Supplementary Material, section 6**).

Then, according to the ANOVA results, a set of contrasts across spatial and temporal conditions was performed by means of planned paired t-tests, for each EEG feature.

Specifically, we assessed a spatial inter-hemispheric contrast for each session (i.e., contrast of values between cortexes, in the pre- and in the post-surgery session; **Figure 2 B, D**), and a pre-post-surgery temporal contrast for each cortex (i.e. contrast of the pre-post difference against 0, in the disconnected and in the contralateral cortex, **Figure 2 C, E**). Further, we ascertained whether the pre-post difference was larger for the disconnected cortex (i.e., contrast of the pre-post difference between cortices; **Figure 2 C, E**); a finding which, in our specific design where measurements were paired over both time and space, would imply that the inter-hemispheric difference changed following surgical disconnection.

**Figure 2.**
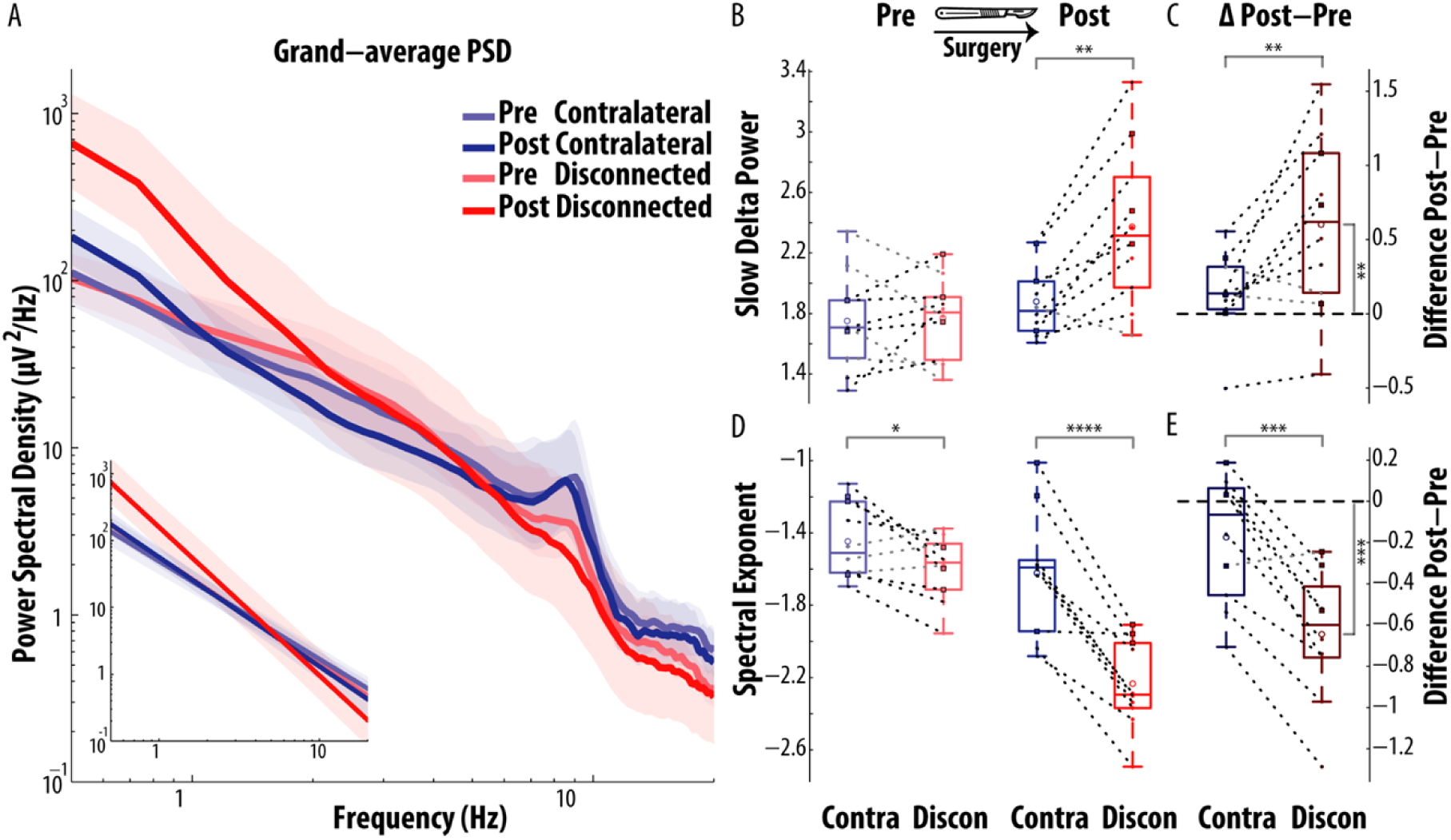
Group-level narrow- and broad-band spectral changes after hemispherotomy reveal a marked EEG slowing of the isolated cortex, robust across patients. **A)** The geometric mean of the PSD was taken first across electrodes, and subsequently across patients, for the observed PSD (main graph) as well as for the aperiodic fit of the PSD (corresponding to a straight line under logarithmic scale for both x and y axes, inset graph). The shaded area indicates the bootstrapped 95% confidence interval for the geometric mean across participants. **B**) The PSD in the Slow-Delta band (0.5-2 Hz) was significantly larger in the disconnected (Discon) than in the contralateral (Contra) cortex, only in the session after surgery. **C)** Following surgery, slow wave activity increased in the disconnected cortex, as indexed by a significant pre to post increase of the Slow-Delta PSD in the disconnected cortex (i.e. positive post-pre differences in the disconnected cortex). This increase was larger in the disconnected than in the contralateral cortex. **D)** The spectral exponent (0.5-20 Hz) was significantly more negative in the disconnected than in the contralateral cortex, in the session before surgery (Pre) and, particularly, in the session after surgery (Post), whereby consistent inter-hemispheric asymmetry was observed in all patients. **E)** Following surgery, the PSD became steeper in the disconnected cortex, as indexed by a significant pre to post decrease in spectral exponent towards more negative values, observed in all patients (i.e. all negative post-pre differences for the disconnected cortex). This decrease was larger in the disconnected than in the contralateral cortex. Patients who received a temporo-parieto-occipital disconnection (squares with a black outline) displayed an overall similar pattern to that of patients who received a hemispherotomy (small dots). Significant contrasts are denoted by star symbols (* P <.05; ** P <.01; *** P <.001; **** P <.0001) over a bracket connecting the contrasted values, either the two cortexes, or the post-pre difference against zero.

We further compared the strength of statistical evidence between narrow-band and broad-band spectral features by means of bootstrap analysis (**Supplementary Material, section 7, Supplementary Figure 4)**.

### Spectral exponent: comparison to reference values during wake and NREM sleep stages

To further establish whether spectral exponent values observed in patients with cortical disconnection were compatible with wakefulness or sleep, we included a reference dataset of spectral exponent values obtained from the EEG of a large sample of neurotypical pediatric participants during wakefulness and sleep (Favaro et al., 2023). In this previous study, the EEG was obtained during a daytime nap opportunity in an outpatient clinic, in children and adolescents aged from 2 to 17 years, who showed no neurological abnormalities. Here, to build the spectral exponent reference dataset, we considered only the subset of participants who were able to fall asleep and reached all NREM sleep stages (up to N3), consisting of 44 participants (24 males, aged between 2-16 years, mean 6.798, s.d.=3.944).

To directly compare spectral exponent values across datasets, we aligned the preprocessing pipeline of the patients with cortical disconnection to that of the previously published study on the reference pediatric sample (Favaro et al., 2023). Hence, in the patients’ dataset, we increased the Butterworth high-pass filter from 0.1 to 0.5 Hz, and consequently restricted the fitting range of the spectral exponent from 0.5-20 Hz to 1-20 Hz (i.e. with the lower bound set at a frequency where the filter effects were dissipated and the expected PSD decay was observed), according to the procedure used in the reference dataset (Favaro et al., 2023). Even after restricting the frequency range under study, the findings on the spectral exponent were consistently replicated in the patient dataset (**Supplementary Material, section 4, Supplementary Figure 3).**

Finally, we adapted the virtual EEG reference of the pediatric reference dataset to that of the patient dataset, which required relatively high spatial specificity. Hence, the spectral exponent of the reference dataset (previously estimated using average reference) was re-estimated over all lateralized nearest-neighbor bipolar derivations (15 per side, 30 total) and the median value was considered. To quantify the proximity of the disconnected and contralateral cortex to reference physiological values of wake and sleep, we estimated the differences of the means of each condition between the two datasets across bootstrap resamplings (**Supplementary Material, section 8).**

### Further comparison of the disconnected hemisphere with physiological NREM Sleep

In the previous part of the study we compared the spectral exponent in patients before and after surgical disconnection to that of a reference pediatric cohort across wakefulness and NREM sleep stages. After observing that the broad-band spectral exponent of the disconnected cortex closely resembled that of deep NREM sleep (of N2 and N3 stages particularly, **Figure 3**), we then explored how the EEG activity of the disconnected cortex differed from physiological sleep, considering NREM sleep stages N2 and N3 together.

**Figure 3.**
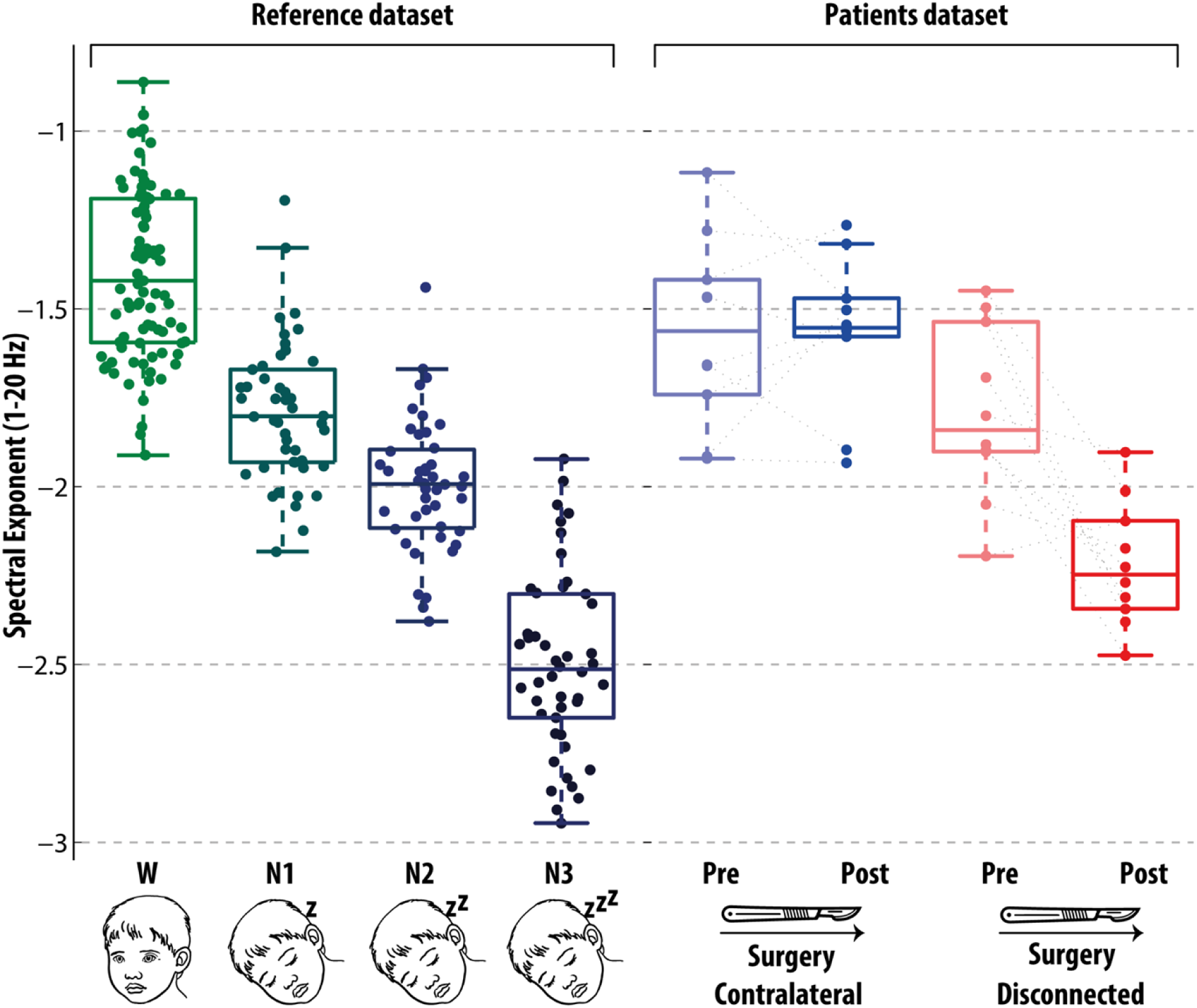
The comparison of the Spectral Exponent in hemispherotomy with a reference pediatric sample across Wake and NREM Sleep reveal values compatible with deep-sleep in the isolated cortex. The spectral exponent is estimated in patients before and after surgery (Patients dataset, right panel), and in a reference pediatric sample across wake and NREM sleep (Reference dataset, left panel). To enable comparison, the spectral exponent is specifically estimated over the 1-20 Hz band, and the cutoff frequency of the pre-processing high-pass filter has been raised to 0.5 Hz, in keeping with Favaro et al., 2023. The contralateral cortex, both before and after surgery, showed values compatible with wakefulness. Before surgery, the pathological cortex showed values intermediate between those of wakefulness and those of N1 sleep. After surgery, the disconnected cortex showed values intermediate between those of N2 and N3 sleep.

This allowed us to ascertain the specific effect of deafferentation of the cortex from subcortical structures, particularly the thalamus, in the formation of two EEG signatures of physiological sleep, namely spindle and slow wave activity. We thus investigated slow wave activity and spindles, and further characterized the underlying spectral and temporal features of the slow-delta and sigma band. We extracted these features in the disconnected cortex, and compared them to those of Wakefulness and NREM sleep of the pediatric reference dataset.

### Slow delta oscillations and slow waves

#### Amplitude of Slow-Delta activity

To compare the strength of slow-delta activity in the disconnected cortex with that of physiological sleep and wakefulness, we estimated the amplitude of the envelope of slow-delta oscillations. For this sake, we first low-pass filtered the EEG signal (IIR forward and reverse Butterworth filter, 5th order) with a cutoff at 2 Hz, in alignment with the spectral domain analysis (**Figure 1-2**), then considered the absolute value of the Hilbert transform. A density plot of the typical span of the amplitude values across subjects is shown in **Supplementary Figure 6**, for both the patient and reference dataset (**Supplementary Material, section 9**). The mean slow-delta amplitude was obtained by averaging the amplitude envelope over time, then its square root was taken, to reduce the skewness of the distribution among subjects; finally, the median value across channels was retained. Further, we corroborated this analysis by means of spectral analysis in the slow-delta band (**Supplementary Material, section 10**).

#### Period of slow delta oscillations and rate of large slow waves

After characterizing the amplitude of slow delta fluctuations, we then characterized the period of slow-delta activity, irrespectively of amplitude, of the disconnected cortex with respect to physiological NREM sleep and wakefulness. To estimate the period of activity showing oscillations in the slow-delta band, the signal was first low-pass filtered in the delta band (IIR forward and reverse Butterworth filter, 5th order, cutoff at 4 Hz), and then only oscillations with a period consistent with slow-delta oscillations was considered (the time between the zero-crossings of a full oscillation was required to be longer than 0.5 sec, corresponding to frequencies lower than 2 Hz, adapted from Molle et al., 2002; Massimini et al., 2004; Piantoni et al., 2013). Differently from mastoid-referenced EEG, where only half-period fluctuations within the negative polarity are considered (Riedner et al., 2007; Bersagliere and Acherman, 2010; Lazar et al., 2015), in our case of bipolar channels we considered full-period signal fluctuations across positive and negative polarity.

We then estimated the mean period across all slow-delta oscillations. A density plot of the typical span of the period values across subjects is shown in **Supplementary Figure 7)**, for both the patient and reference dataset. To compare values between subjects of different conditions, the median value across channels was retained (**Figure 4**).

**Figure 4.**
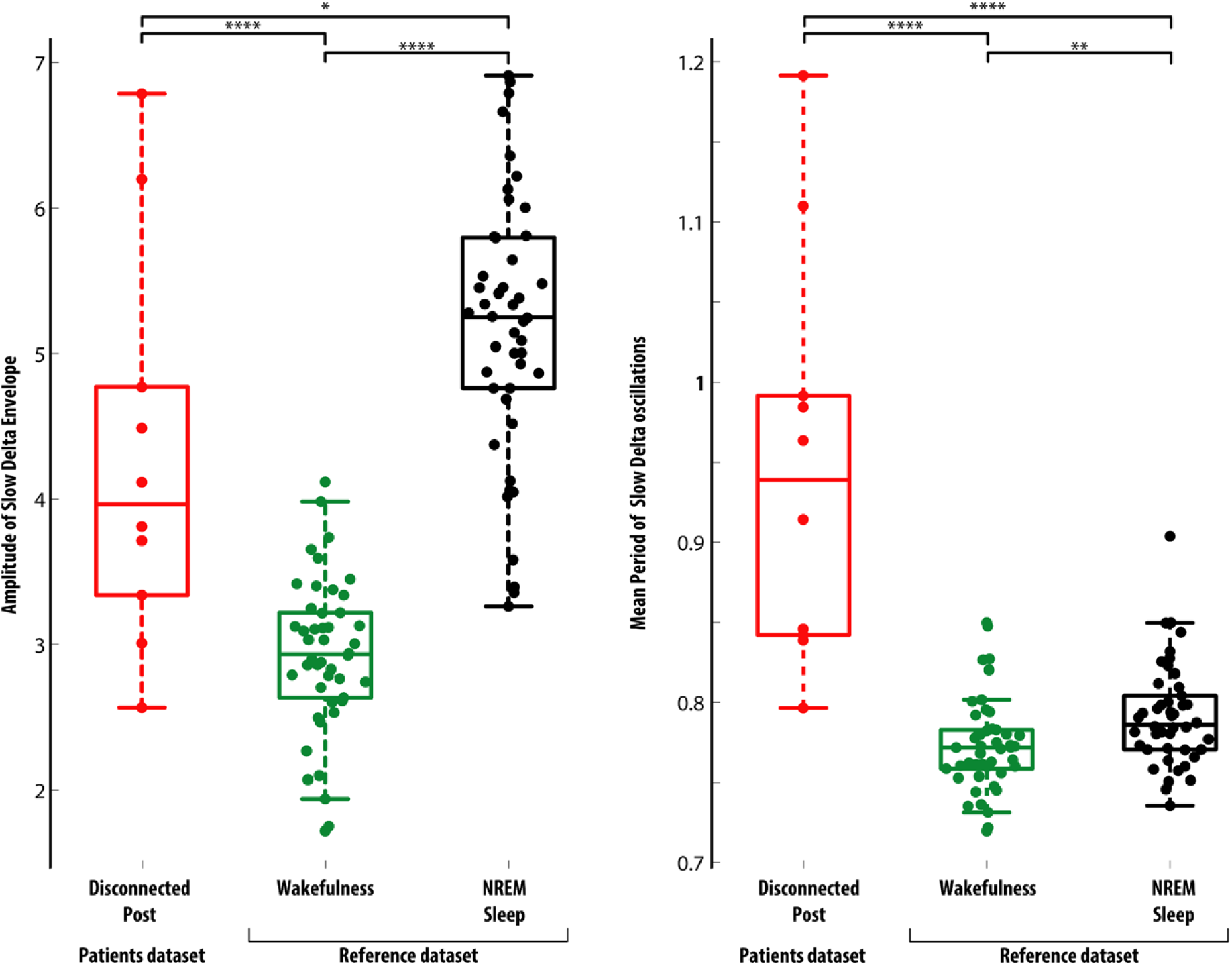
Mean amplitude of the slow-delta envelope and period of slow-delta oscillations in the disconnected hemisphere, compared with respect to wakefulness and NREM sleep in the reference pediatric sample. A) The mean amplitude of the Envelope filtered in the slow-delta band, median across channels, is shown for each patient in the disconnected cortex after surgery, and, for comparison, in the reference dataset during wakefulness and NREM sleep. B) The mean period of slow-delta oscillations, median across channels, is shown for each subject across conditions. The median across channels is shown for each subject across conditions.

#### Sigma oscillations and spindle activity

To determine whether the disconnected cortex (recorded when the patients were behaviourally awake) was devoid of spindle oscillations, a trained neurologist first inspected the EEG traces. Despite the abundance of slow oscillations resembling a sleep-like pattern, spindle grapho-elements were not identified and only sporadic, low-amplitude alpha activity could be identified in the disconnected cortex, in line with the established thalamic origin of sleep spindles (Luthi, 2014; Fernandez and Luthi, 2020). Subsequently, quantitative spectral and temporal analyses were devoted to confirm this qualitative finding.

#### Power of Sigma Periodic activity

We aimed to estimate the portion of the PSD in the sigma band generated by spindle activity, knowing that a strongly rhythmic signal contributes to a distinct narrow-band PSD peak, that broad-band changes can bias narrow-band PSD estimates, and finally, taking into account that the sigma band overlaps with the alpha and beta bands.

Specifically, changes in broad band activity could differently affect the estimates of sigma power between conditions: the disconnected cortex displayed a notable broad-band rotation of the PSD and increase in low-frequency power following surgical disconnection (similarly to what was observed in the physiological wake-sleep transition, **Figure 5, panel A, B**), thus affecting the amount of narrow-band total power in the sigma band.

**Figure 5.**
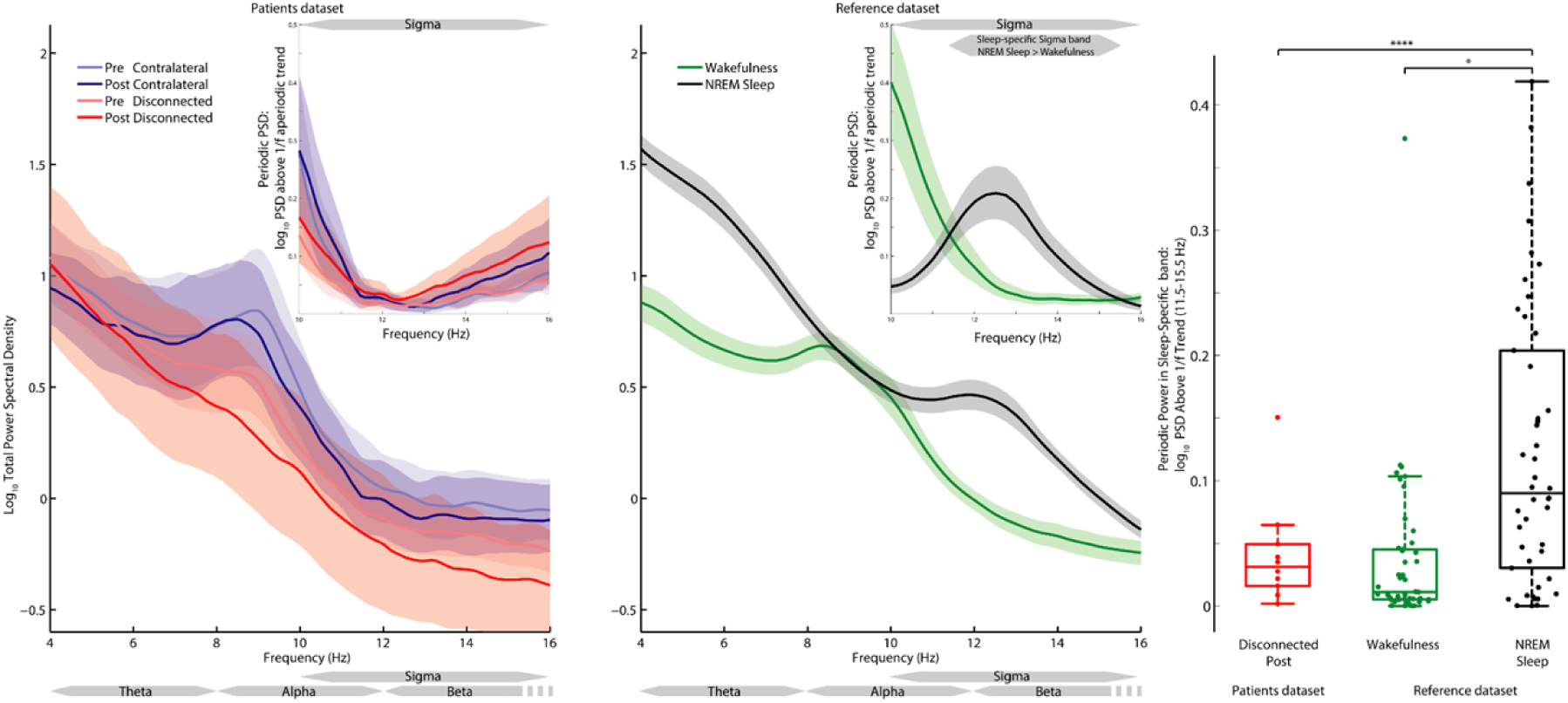
Analysis of total and periodic PSD in the sigma band reveal the absence of rhythmic spindle activity in the disconnected cortex. **A)** Grand-average PSD in the patient dataset is shown for the disconnected and contralateral cortex, before and after surgery. A broad range of frequencies (4-16 Hz) spanning theta, alpha, and the lower range of the beta band is shown to provide a context for the PSD in the sigma band. The median values across channels is obtained, then the mean values across subjects is shown, along with its bootstrapped confidence interval. The inset displays the periodic PSD component, i.e. the PSD exceeding above the 1/f-like trend (displayed in Figure 2), within the sigma band (10-16 Hz). While the disconnected cortex displayed predominantly broad-band aperiodic activity with no clear peaks in the total PSD, its periodic PSD component displayed residual power pertaining to the neighboring alpha and beta bands; however no sigma peak occurred within the sigma band. **B)** Grand-average PSD is shown for physiological Wakefulness and NREM sleep (stages N2 and N3 combined) obtained from the pediatric cohort of the reference dataset. In the inset, the periodic PSD component of Wakefulness displays a downward inflection from the alpha band, while that of sleep displays a clear peak centered in the sigma band–reflecting spindle rhythmic activity. A thick grey line highlights the sleep-specific sigma band (11.5-15.5 Hz), where the grand-average periodic sigma PSD of NREM Sleep exceeds that of Wakefulness. **C)** The mean periodic component of the PSD in the sleep-specific sigma band (11.5-15.5 Hz), median across channels, is shown for each subject. The disconnected cortex showed values overlapping and consistent with those of Wakefulness, and substantially lower than those of NREM sleep.

Since the PSD of the EEG signal originates from a mixture of aperiodic and periodic activity, it can be accordingly decomposed as a sum of aperiodic and periodic PSD components (Palva and Palva, 2018; Donoghue et al., 2022), by taking into account that the aperiodic PSD has a 1/f-like shape. Thus, to estimate the power of sigma activity corresponding to genuinely periodic activity (i.e. spindles), we considered only the power that exceeded the 1/f-like aperiodic trend (i.e. the positive PSD residuals after subtracting the 1/f-trend from the total PSD, **Figure 5, insets of panel A, B**).

During physiological sleep, the PSD displays a clear peak in the sigma band, a hallmark of the presence of spindles (**Figure 5, panel B**). Yet, the sigma band (∼11-15 Hz, see Purcell et al., 2017) overlaps with both the alpha (∼ 8-12 Hz) and beta band (∼ 12-30 Hz) (**Figure 5, inset of panel B**). Thus, we restricted the sigma frequency range of interest to a band where the grand-average periodic PSD of sleep exceeded that of wakefulness in the reference sample, to obtain a more sleep-specific estimate of sigma power, with reduced influence from wake-like activity. We thus estimated the amount of periodic sigma power by averaging the PSD in a sleep-specific sigma band, as determined from the overall sleep-wake difference (11.5-15.5 Hz, **Figure 5, inset of panel B)**. The strength of sigma oscillations – as defined above – in the disconnected cortex was then compared to that of sleep and wakefulness in the reference sample. We expected the disconnected cortex would show values considerably lower than those of sleep, and compatible with (or lower than) those of wakefulness.

Moreover, in the **Supplementary Material, section 11** we detected bursts of sigma oscillations, following a time-domain approach commonly employed in spindle detection during sleep (Ferrarelli et al., 2007; Andrillon et al., 2011; Sarasso et al., 2014; Hahn et al., 2020). If the properties of sigma bursts (such as their rate and amplitude) in the disconnected cortex would differ significantly from those observed during physiological sleep and resemble those seen during physiological wakefulness instead – where spindles are notably absent – this would further support the notion that spindles are also absent in the disconnected cortex.

## Results

### Patients and EEG recordings

Among the 10 patients fulfilling inclusion criteria, 4 had a diagnosis of focal drug-resistant epilepsy with structural etiology on a malformative basis (i.e. pachygyria, polymicrogyria or focal cortical dysplasia) and 6 as an outcome of perinatal hypoxic-ischemic insult. Patients underwent a complete disconnection of a large cortical portion: either an entire hemisphere (complete hemispherotomy, 7 patients), or the temporo-parieto-occipital region (3 patients). The EEG recordings were performed within 12 months before surgery (mean=5.6, std = 4.67, range = [1, 12]) and at least six months after surgery (delay in months: mean=19.8, std=9.6, range = [6.96, 34.92]). The age at pre-surgery EEG spanned from 2 to 11 years (mean = 7.8, std = 2.9); in each recording we identified at least 3 min of quiet wakefulness free from artifacts and interictal epileptiform discharges (mean= 7.379, std= 3.489 range = [3.561, 12.815]). Demographic and clinical information for each patient are reported in **Supplementary Table 3.**

### Surgical disconnection and EEG slowing

A representative example of the main results is shown for an 11-year-old girl with a left porencephalic lesion secondary to perinatal stroke undergoing hemispherotomy (**Figure 1A**). The visual inspection of the EEG time-series and of the topographies of low-frequency PSD (Slow-Delta power, 0.5-2 Hz) reveal high-amplitude (>50 microvolts) slow waves emerging over the disconnected cortex post-surgery, associated with pronounced inter-hemispheric asymmetry (**Figure 1B)**. Further, the PSD profiles (**Figure 1C**) indicate that the narrow band increase in the Slow-Delta band was accompanied by an overall redistribution of power from high to low frequencies, resulting in a steeper broad-band decay of the disconnected cortex following surgery. These observations were quantified by the Slow-Delta power and the spectral exponent (indexing the slope of the PSD decay over the 0.5-20 Hz range) over intra-hemispheric bipolar derivations (**Figure 1D**).

These single-subject findings were consistently observed at the whole sample level. When comparing the seven patients with complete hemispherotomy to those who underwent temporo-parieto-occipital disconnection, we observed a highly similar pattern of localized increase in the degree of EEG slowing in the disconnected cortex (**Supplementary Material, section 5**). Results are thus reported across all 10 patients, irrespective of the extent of the surgery. As shown in **Figure 2**, at the population level the disconnected cortex following surgery displayed increased Slow-Delta power and an overall PSD redistribution towards low frequencies and a steeper broad-band decay, as evident from the PSD and its 1/f-like fit, aggregated across patients. These observations were confirmed by statistical analysis, reported in the next section.

### Asymmetry of cortical states after disconnection

For each of the two EEG dependent variables (Slow Delta Power and Spectral Exponent, indexing narrow-band and broad-band slowing respectively), the ANOVA on the mixed effects model revealed a main effect of time (both p <0.001), demonstrating a change due to surgical disconnection. Further, it also revealed a significant interaction effect between time and space (both p <0.001), indicating that surgery differently affected the disconnected vs the contralateral cortex, and, similarly, that the inter-hemispheric asymmetry changed following surgery (for details see **Supplementary Material, section 6).**

The specific effects of cortical disconnection on each EEG feature were explored through a series of planned pairwise contrasts, t-tests with 9 d.f. (**Figure 3, Supplementary Table 2**).

Contrasting the pre-post difference between cortices, the disconnected cortex underwent a greater change following surgery as compared to the contralateral, for both the Slow-Delta power (T(9)= -3.146, p=0.0118, greater change observed in 8/10 patients, **Figure 2 C**) and the spectral exponent (T=6.464, p=0.0001, greater change observed in 9/10 patients, **Figure 2 E**), indicating also that the inter-hemispheric asymmetry increased following surgery.

When assessing spatial inter-hemispheric contrasts (contralateral vs disconnected cortex), we found that Slow-Delta power was significantly larger in the pathological cortex than in the contralateral cortex, but only in the post-surgery session (Pre: T=-0.228, p=0.8246; larger in 6/10 patients; Post: T = -4.245, p=0.0022, larger in 9/10 patients, **Figure 2B**); the spectral exponent was significantly more negative in the disconnected cortex than in the contralateral, both pre-surgery (T=2.395, p=0.0402; more negative in 7/10 patients) and, especially, post-surgery (T=7.118, p<0.0001, more negative in 10/10 patients, **Figure 2D**).

Assessing temporal contrasts (Post-Pre surgery), the Slow-Delta power was larger following surgery, significantly only in the disconnected cortex (contralateral: T= -1.462, p=0.1777, larger in 9/10 patients; disconnected: T= -3.215, p=0.0106; larger in 10/10 patients, **Figure 2C**); similarly, the spectral exponent was significantly more negative following surgery, only in the disconnected cortex (contralateral: T= 1.786, p=0.1077, more negative in 5/10 patients; disconnected: T=6.455, p=0.0001, more negative in 10/10 patients, **Figure 2E**).

Overall, before surgery we found no evidence for interhemispheric difference in narrow-band Slow-Delta power and only a small, albeit significant, inter-hemispheric asymmetry in the degree of broad-band slowing. Instead, after surgery, we observed a marked inter-hemispheric asymmetry characterized by prominent increase in both the degree of EEG narrow-band and, especially, of broad-band slowing over the disconnected cortex. The spectral exponent revealed larger effect sizes than delta-power, thus potentially providing a more sensitive index of the neurophysiological consequences of hemispherotomy (**Supplementary Material, section 7** and **Supplementary Figure 4**).

### The awake spectral exponent of the disconnected cortex aligns with values of physiological NREM sleep

Finally, the spectral exponent values assessed in patients before and after cortical disconnection were compared to the values assessed in a reference pediatric sample during wakefulness and NREM sleep (**Figure 3**). Qualitatively, the typical values (i.e., those of the central half of the sample, within the interquartile range) of the contralateral cortex overlapped with the typical values observed during wakefulness and N1 in the reference sample. The typical values of the disconnected cortex before surgery overlapped with the typical wakefulness and N1 values of the reference sample; after surgery, the typical values of the disconnected cortex overlapped with the typical N2 and N3 values of the reference sample. Notably, 9/10 values from the disconnected cortex did not overlap with the wakefulness distribution; and the 10th case was below all but one of the values from the wakefulness distribution.

Subsequently, we compared the mean spectral exponent values of each condition across the two datasets through bootstrap resampling. Specifically, we considered the bootstrapped distribution of the pairwise differences between the mean values of the pediatric reference participants (either during wakefulness, N1, N2, or N3 sleep) and the mean values of the patients (in the disconnected or contralateral cortex, before or after surgery) (**Supplementary Material, section 8; Supplementary Figure 5**). According to bootstrapped differences of mean values, the homologous contralateral cortex, both before and after surgery, showed values distinct from the NREM sleep distributions and closest to the wakefulness distribution, albeit on the lower end. Moreover, the disconnected cortex before surgery showed values similar to the N1 sleep distribution; crucially, after surgery the disconnected cortex showed values distinct from the wakefulness distribution and intermediate between those of the N2 and N3 sleep distributions.

### Slow-Delta activity in the disconnected cortex

We further characterized slow wave activity occurring during wakefulness in the disconnected cortex, by comparing the average amplitude of the envelope of EEG activity filtered in the slow-delta band with respect to the pediatric reference sample across both wake and sleep. Confirming the results of spectral domain analysis, large amplitude values occurred more frequently in the pathological cortex following surgical disconnection, as displayed in the grand-average density plot **(Supplementary Figure 6).** Most notably, larger amplitude values were found with respect to the awake pediatric reference sample (T(54)=−5.3132, P=2.0934e−06). When directly compared to physiological N2 and N3 sleep, slow wave activity appearing during wakefulness in the disconnected cortex was on average smaller (T(54)=2.5986, P=0.012043). Similar findings were obtained considering the PSD in the slow-delta band **(Supplementary Material, section 10)**

Next, we characterized the typical period of slow-delta oscillations by analysing activity in a wider delta band and considered only slow-delta oscillations (period >0.5 s). Large period values occurred more frequently in the pathological cortex following surgical disconnection, as displayed in the grand-average density plot **(Supplementary Figure 7).** The mean period of slow-delta oscillations of the disconnected cortex was markedly larger than that observed during either wakefulness (T(54)=−8.566, P=1.2167e−11) or sleep (T(54)=−7.5059, P=6.2164e−10). Overall, the disconnected cortex showed a remarkable emergence of slow-delta activity, leading to a state departing from wakefulness, yet different from full-fledged NREM sleep.

### Absence of spindles in the disconnected cortex

During physiological NREM sleep, spindle activity, due to its narrow-band oscillatory property, is marked by a pronounced PSD peak in the sigma frequency band. The PSD of the disconnected cortex did not display any peak in the sigma band, similarly to the reference sample during wakefulness (**Figure 5**, panel A, B).

We then extracted the periodic PSD component, by considering only the PSD that exceeded the 1/f-like aperiodic trend (shown in **Figure 1**, inset of panel A, **Figure 2**, inset of panel B), to better disentangle signs of periodic activity in the sigma frequency band generated by potential spindles. In the disconnected cortex, the periodic PSD component (i.e. that exceeding the aperiodic 1/f background) in the 10-16 Hz band originated from the upper-end of the alpha band and from the lower end of the beta band **(Figure 5, inset of panel A)**. The presence of genuine periodic activity in the sigma band, as marked by a distinct sigma peak, could thus be ruled out at the group level. This was corroborated by the inspection of individual periodic PSD components.

To obtain a marker specific for sleep-like sigma activity, we subsequently considered the portion of the sigma band that was specific for sleep spindles, i.e. whereby the periodic PSD in the sigma band of sleep exceeded that of wakefulness. The grand-average periodic sigma power of NREM sleep exceeded that of wakefulness in the 11.5-15.5 Hz range in the reference dataset (**Figure 5, inset of panel B**); we accordingly considered the mean periodic sigma power in this sleep-specific sigma band. In the disconnected cortex, the mean periodic power in the sleep-specific sigma band was remarkably lower than that of NREM sleep and similar to that observed during physiological wakefulness. Hence, spindle activity, clearly marked by pronounced periodic sigma power during physiological sleep, was absent in the disconnected cortex, similarly to what was observed during physiological wakefulness.

Overall, the absence of any sleep-like sigma peak in the PSD **(Figure 5)**, concurrently with the presence of occasional bursts of energy in the sigma band similar to those of wakefulness (**Supplementary Material, section 11**), showed that the disconnected cortex did not display any sleep spindles. All the above quantitative analyses confirmed the visual assessment made by the neurologist.

## Discussion

This study represents the first investigation of the electrophysiological changes occurring in awake humans after the surgical disconnection of extensive cortical regions, including an entire hemisphere. Unlike the contralateral homologous cortex, after surgical disconnection the isolated cortex entered a state dominated by large-amplitude slow wave activity (below 2 Hz), and an overall redistribution of the PSD from high to low frequencies. Such broad-band neurophysiological slowing, indexed by the spectral exponent, was similar to that previously found in the physiological (Shen et al., 2003; Miskovic et al., 2019; Bódizs et al., 2021; Zilio et al., 2021; Schneider et al., 2022; Horváth et al., 2022; Alnes et al., 2023; Rosenblum et al., 2024; Bódizs et al., 2024), pharmacological (Colombo et al., 2019; Maschke et al., 2023a; Maschke et al., 2023b) and pathological unconscious states (Colombo et al., 2023; Maschke et al., 2023a; Zilio et al., 2023). Upon direct comparison with a reference pediatric sample, the disconnected cortex showed a broad-band spectral exponent resembling deep NREM sleep. Yet, the isolated cortex showed distinctive features compatible with a lack of thalamic modulation, notably the absence of spindle, and slow-delta activity with smaller amplitude and prolonged period with respect to NREM sleep.

### Mechanisms of EEG slowing in cortical disconnection

Following surgery, prominent slow waves with a pronounced increase in Slow-Delta power (0.5 - 2 Hz) appeared over the disconnected cortex. The contralateral cortex, retaining intact subcortical inputs, did not display any significant increase in the degree of EEG slowing following surgery, despite the surgical resection of all its inter-hemispheric connections. This finding, in line with neurophysiological evidence in callosotomy (Quattrini et al., 1997; Pizoli et al., 2011), suggests that the primary cause of EEG slowing observed in the disconnected cortex lies in the deafferentation from subcortical, rather than cortical, inputs.

During NREM sleep, EEG slow waves emerge throughout the cerebral cortex as a consequence of decreased levels of activating neuromodulation from brainstem activating systems (McCormick and Williamson, 1989). A similar pattern can be observed in pathological conditions following lesions or compressions in the brainstem and midbrain ascending activating systems (McCormick 1992; Lee and Dan Y, 2012). More generally, slow waves can be found in various experimental models in which cortical circuits are disconnected from subcortical inputs, such as in isolated cortical gyri (i.e., cortical slabs) (Timofeev et al., 2000), in cortical slices in vitro (Capone et al., 2019) as well as after thalamic inactivation (Lemieux et al., 2014). As such, cortical slow waves are considered the elemental intrinsic regime of cortical circuits (Sanchez-Vives et al., 2017). The present study in hemispherotomy provides further evidence for the account of cortical slow waves as the default activity pattern of the cortical network (Sanchez-Vives et al., 2017). Remarkably, this is, to our knowledge, the first evidence that this pattern can persist for months and years (recordings were performed between 6 and 36 months post surgery) after complete cortical disconnection.

Below, we discuss the significance of these findings in relation to the ongoing debate about the potential for “islands of awareness” following hemispherotomy and in the broader perspective of the consequences of structural brain disconnections.

### The problem of inaccessible awareness

Inferring consciousness in isolated human cortical hemispheres represents a unique challenge (Bayne et al., 2020; Gauvry and Rüber, 2024). For example, the surgically disconnected cortex differs from lab-grown cortical organoids, because it has developed as part of a human brain in contact with the external world, and thus some arguments against the possibility of consciousness in cortical organoids would not apply to hemispherotomy (Croxford and Bayne, 2024).

A vast literature on acquired brain-injury (Gschaidmeier et al., 2021; Asaridou et al., 2020; Sebastianelli et al., 2017), hemispherectomy (Dandy, 1928; Crockett and Estridge, 1951; Peacock et al., 1996; Lew, 2014; Fisher et al., 2022) and split-brain patients (Schechter and Bayne, 2021; de Haan et al., 2018) suggests that one hemisphere is sufficient to support consciousness. However, hemispherotomy also differs from split-brain surgery, where the two hemispheres are disconnected from each other but retain all subcortical afferents and their ability to interact with the external world (Schechter and Bayne, 2021; de Haan et al., 2018). Hence, whether the isolated cortex represents a distinct locus of consciousness from that supported by the contralateral hemisphere remains an open question. Two recent fMRI studies indicate preserved neural networks lateralized within the isolated hemisphere and specifically the integrity of the default mode network (DMN) (Blauwblomme et al. 2020; Rüber et al., 2024). While the first study (Blauwblomme et al. 2020) interprets this result as indicative of a lower level of consciousness, the second (Rüber et al., 2024) contemplates the possibility of an island of awareness. A possible rationale for the second interpretation is that, in healthy awake participants, the DMN is typically activated during self-directed mentation, such as daydreaming and reflection (Menon, 2023). Yet, resting state networks, including the DMN, can be disrupted in conscious psychedelic states (Gattuso et al., 2023) and, crucially, preserved in unconscious states such as sleep and anesthesia (Boly et al., 2008; Horovitz et al., 2008; Vincent et al., 2007; Akeju et al., 2014) making it difficult to infer consciousness solely based on fMRI patterns.

The present findings of prominent slow waves and steep spectral decay strongly suggest that the disconnected cortex rests in an electrophysiological state that is not compatible with the presence of an island of awareness. This conclusion is supported by different lines of converging evidence.

First, EEG slow waves are canonically associated with unconscious conditions encompassing NREM sleep (Jouvet, 1967; Nir et al., 2011; Pigorini et al., 2015), general anesthesia (Brown et al., 2010; Purdon et al., 2015; Murphy et al., 2011; Lewis et al., 2012; Sarasso et al., 2015) and the vegetative state (Comanducci et al., 2020; Colombo et al., 2023; Estraneo et al., 2016; Schiff, 2016; Lechinger et al., 2013; Rosanova et al., 2018). As shown by animal and human studies, EEG slow waves reflect the tendency of cortical neurons to become silent after an initial activation, a mechanism that prevents cortical circuits from engaging in the complex network interactions normally observed in conscious states (Pigorini et al., 2015; Rosanova et al., 2018; D’Andola et al., 2018; Arena et al., 2021).

Second, recent studies in healthy humans have shown that increased low frequency power and decreased high frequency power are predictive of the absence of dream reports during sleep (Siclari et al., 2017). Therefore, the present finding of a redistribution of power from high to low frequencies detected by the spectral exponent (**Figure 1** and **Figure 2**) can be interpreted as an indication of the reduced likelihood of dream-like experiences in the isolated cortex.

Third, the spectral exponent has been previously tested as an index of consciousness in large samples of healthy controls and patients, including challenging conditions of sensorimotor disconnection and behavioral unresponsiveness. In good agreement with perturbational measures of brain complexity (the Perturbational Complexity Index, see Casali et al., 2013), the spectral exponent robustly discriminated between conscious and unconscious states, across wakefulness, sleep, physiological or pharmacological dreams, anesthesia induced by different compounds, and severe brain injury (Colombo et al., 2019; Colombo et al., 2023; Maschke et al., 2023b; Casey et al., 2023). Overall, detecting an electrophysiological pattern aligned with that observed in reference physiological, pharmacological and pathological conditions in which the global state of consciousness is abolished or greatly reduced, provide evidence against the possibility that hemispherotomy may result in cortical islands of awareness.

More broadly, the present findings prompt further investigation into the relationship between EEG slowing and fMRI functional connectivity patterns. While macroscale fMRI resting-state networks, including the DMN, remain preserved in the isolated hemisphere, intrahemispheric connectivity increases, and the normal anticorrelations between networks are lost. Recent rodent studies suggest a link between functional network abnormalities, such as DMN hyperconnectivity, and EEG slow waves (Rocchi et al., 2022). Similarly, stroke patients show EEG slowing and altered connectivity patterns, including the loss of normal anticorrelations, reduced interhemispheric connectivity, and increased intrahemispheric connectivity (Massimini et al. 2024). In this context, the unilateral sleep-like state observed after hemispherotomy offers a unique model for exploring the interplay between electrophysiological dynamics and fMRI functional networks across both physiological and pathological brain states.

### Disconnection and sleep-like dynamics

The generation of sleep-like slow waves following structural injuries and disconnections in awake individuals is a long-standing concept. In fact, the term delta waves—now commonly associated with the EEG patterns of deep NREM sleep—was originally introduced nearly a century ago by William Grey Walter to describe the slow waves recorded during wakefulness over focal brain lesions (Walter, 1937). The idea that cortical sleep-like dynamics can intrude during wakefulness after brain injury, leading to network disruptions, loss of consciousness, or cognitive and motor deficits, has recently been systematically revisited (Massimini et al., 2024). Delta waves during wakefulness have been observed in various conditions, such as traumatic brain injury, diffuse axonal injury, and ischemic or hemorrhagic stroke (Gloor et al., 1977; Meythaler et al., 2001; Sarasso et al., 2020; Lanzone et al., 2022; Massimini et al., 2024). However, their relationship to the key neurophysiological features of physiological sleep has yet to be systematically assessed.

The surgical procedure of hemispherotomy offers a unique model for investigating and characterizing the intrusion of sleep-like slow wave activity during wakefulness following structural damage. In this study, we leveraged this model by directly comparing EEG recordings from hemispherotomy patients with NREM sleep recordings in a pediatric reference dataset, which spans the typical age range of patients undergoing cortical disconnection (Favaro et al., 2023).

First, we assessed the overall degree of broadband EEG slowing using the spectral exponent. Following surgery, the spectral exponent of the isolated cortex deviated from the wakefulness distribution observed in the pediatric sample in 9 out of 10 cases (Figure 3)—a finding consistent with N2-N3 sleep, as confirmed by bootstrap analysis (see **Supplementary Figure 5**). However, the EEG pattern recorded after disconnection also exhibited key differences from that of physiological sleep.

Although dominated by slow-wave activity and exhibiting a broadband spectral decay comparable to that found in the sleeping reference sample, the isolated cortex displayed slow-delta activity with smaller amplitude and longer periods compared to that seen during physiological sleep (Figure 5). These differences are likely primarily due to thalamic disconnection. The integrity of cortico-thalamo-cortical loops is known to enhance the synchronization—and consequently the amplitude—of EEG slow oscillations. Additionally, thalamic neurons play a role in initiating cortical up-states (David et al., 2013), and focal pharmacological inactivation of the thalamus has been shown to extend the period of cortical slow waves (Lemieux et al., 2014) and prolong the duration of down-states (Zucca et al., 2019). Beyond the effects of deafferentation from ascending thalamic input, diminished lateral excitatory connectivity may also contribute. In cortical slabs, the period of slow oscillations has been found to scale inversely with the size of the isolated gray matter (Timofeev et al., 2000), suggesting that reduced recurrent lateral excitation results in slower recruitment of cortical activity and less frequent up-states (Lemieux et al., 2014).

Crucially, the disconnected cortex showed the emergence of slow waves that were not accompanied by spindle activity. Analysis of sigma-band activity revealed the absence of a peak in both the total PSD and in the periodic PSD component, consistent with the characteristics of physiological wakefulness (Figure 4, Cellier et al., 2021; Favaro et al., 2023). This observation was further supported by the analysis of sigma bursts (**Supplementary Material, section 11**). This result is unsurprising, as spindles are generated in the thalamus and require intact bidirectional thalamo-cortical connectivity (David et al., 2013), but raises interesting questions about the role of slow waves after disconnection.

### Limitations and future directions

Even before surgery, the pathological cortex exhibited slightly lower mean spectral exponent values compared to its contralateral counterpart, with a distribution centered between those of wakefulness and N1 sleep in the reference pediatric sample (Figure 3). This relative slowing under baseline conditions may be influenced by interictal epileptiform discharges, although these pathological grapho-elements were meticulously excluded during data pre-processing by a trained neurologist (see **Supplementary Figure 1**). Additional contributing factors could include focal cortical malformations and gliosis (Panet-Raymond and Gotman, 1990; Vanrumste et al., 2005; Blümcke et al., 2011). Alternatively, some degree of slowing may be contributed by local slow waves that play a protective role against aberrant epileptiform activity (Sheybani et al., 2023). In this context, the role of the pathological hemisphere and its contribution to consciousness prior to surgery remain open questions. While the current study used available retrospective clinical data before and after hemispherotomy surgery, future prospective studies should acquire EEG at standardized intervals.

The main finding of this study concerns the relative changes observed in the disconnected cortex after surgery compared to the contralateral hemisphere. The broadband slowing, as indexed by the spectral exponent, aligns with patterns observed in unconscious reference conditions, alleviating concerns about the presence of an “island of awareness” within the disconnected hemisphere. However, a general caveat of this conclusion is that any inference about the presence or absence of consciousness, based solely on the brain’s physical properties (including spontaneous EEG dynamics), must be approached with caution, particularly in neural structures that are not behaviorally accessible. It remains possible that some form of consciousness could persist even in the presence of prominent slow waves in the spontaneous scalp EEG, as revealed by peculiar genetic conditions, i.e. Angelman syndrome (Frohlich et al., 2021), and by transient, disconnected psychedelic states (Blackburne et al., 2024) . This possibility warrants further exploration. High-density EEG recordings of the disconnected hemisphere could help confirm the pervasiveness of sleep-like slow waves beyond the coarse spatial resolution of standard EEG. Additionally, perturbational approaches that combine transcranial magnetic stimulation (TMS) and EEG (Casali et al, 2013; Casarotto et al., 2016), which are highly sensitive to detecting consciousness even in the presence of strong delta activity (Comanducci et al., 2023; Casarotto et al. 2024), may provide further insights. Furthermore, in cases where olfactory pathways to the isolated cortex remain intact, it will be important to assess whether higher-level processes, such as the discrimination between familiar and unfamiliar odors, are preserved (Gauvry and Rüber, 2024).

Finally, the discovery of a cortical sleep-like state persisting for years in the absence of the subcortical modulation characteristic of physiological sleep raises significant questions. Specifically, the functional significance of cortical slow waves intruding during wakefulness— in absence of the subcortical inputs required for spindle activity—remains unclear. Given that similar sleep-like dynamics are commonly observed following a wide range of brain injuries (Massimini et al., 2024), it is crucial to determine whether these patterns merely represent a regression of cortical activity to its default state (Krone and Vyazovskiy, 2021) or whether they serve plastic or homeostatic functions (Rosanova et al., 2018; Tscherpel et al., 2020; Bai et al., 2022; Sarasso et al., 2024; Harquel et al., 2024; Tscherpel et al., 2024). Addressing this question is particularly important for understanding the role of slow waves after brain injury, their potential implications for rehabilitation, and will require focused longitudinal investigations.

## Data availability

The original EEG and imaging data are sensitive, as they pertain to pediatric patients. These data can only be made available on a public repository in aggregated form.

The Matlab code and an equivalent Python code translation are available online https://github.com/milecombo/spectralExponent/blob/master/README.md.

## Acknowledgements

Thanks to all the children and their parents who made this work possible. Thanks to the hospital technicians who performed the standard EEG recordings. Jacopo Favaro thanks the TMO group for inspiring discussions and the constant support. Thanks to Gianluca Gaglioti for valuable insights and discussions.

## Glossary

aperiodic PSD component: the portion of the PSD with a 1/f-like shape
ANOVA: Analysis of Variance
Contra: contralateral cortex, homologous of the disconnected
Discon: Disconnected cortex
DMN: Default Mode Network
d.f.: Degrees of Freedom for statistical tests
EEG: Electroencephalography
EMG: Electromyography
EOG: Electrooculography
fMRI: Functional Magnetic Resonance Imaging
Hemispherotomy: the disconnection of a large portion of the cortex, up to an entire hemisphere
‘island of awareness’: a cortical area disconnected from the environment yet capable of awareness
IIR: Infinite Impulse Response, type of filter
ICA: Independent Component Analysis
MRI: Magnetic Resonance Imaging
MCS: Minimally Conscious State, diagnosis in the spectrum of Disorders of Consciousness
N1: Stage 1 Non-Rapid Eye Movement Sleep
N2: Stage 2 Non-Rapid Eye Movement Sleep
N3: Stage 3 Non-Rapid Eye Movement Sleep
NREM: Non-Rapid Eye Movement
periodic PSD component: the portion of the PSD exceeding the 1/f-like decay
Pre: Pre hemispherotomy surgery
Post: Post hemispherotomy surgery
PSD: Power Spectral Density, estimates of the power density across the frequency spectrum
Slow-Delta power: Mean PSD between 0.5-2 Hz
Spectral exponent: reflects the decay of the aperiodic PSD component, i.e. the degree of broad-band slowing
Spindle grapho-elements: Rhythmic bursts of EEG activity (∼10-16 Hz in pediatric population) during NREM sleep
TMS: Transcranial Magnetic Stimulation
1/f: One over f, the aperiodic PSD component is inversely proportional to the frequency (f)

